# Reward Modulates Local Field Potentials, Spiking Activity and Spike-Field Coherence in the Primary Motor Cortex

**DOI:** 10.1101/471151

**Authors:** Junmo An, Taruna Yadav, John P. Hessburg, Joseph T. Francis

## Abstract

Reward modulation of the primary motor cortex (M1) could be exploited in developing an autonomously updating brain-machine interface (BMI) based on a reinforcement learning architecture. In order to understand the multifaceted effects of reward on M1 activity, we investigated how neural spiking, oscillatory activities and their functional interactions are modulated by conditioned stimuli related reward expectation. To do so, local field potentials (LFPs) and singleunit/multi-unit activities were recorded simultaneously and bilaterally from M1 cortices while five non-human primates performed cued center-out reaching or grip force tasks either manually using their right arm/hand or observed passively. We found that reward expectation influenced the strength of alpha (8-14 Hz) power, alpha-gamma comodulation, alpha spike-field coherence, and firing rates in general in M1. Furthermore, we found that an increase in alpha-band power was correlated with a decrease in neural spiking activity, that firing rates were highest at the trough of the alpha-band cycle and lowest at the peak of its cycle. These findings imply that alpha oscillations modulated by reward expectation have an influence on spike firing rate and spike timing during both reaching and grasping tasks in M1. These LFP, spike, and spike-field interactions could be used to follow the M1 neural state in order to enhance BMI decoding (An et al., 2018; Zhao et al., 2018).

**Significance Statement:** Knowing the subjective value of performed or observed actions is valuable feedback that can be used to improve the performance of an autonomously updating brain-machine interface (BMI). Reward-related information in the primary motor cortex (M1) may be crucial for more stable and robust BMI decoding (Zhao et al., 2018). Here, we present how expectation of reward during motor tasks, or simple observation, is represented by increased spike firing rates in conjunction with decreased alpha (8-14 Hz) oscillatory power, alpha-gamma comodulation, and alpha spike-field coherence, as compared to non-rewarding trials. Moreover, a phasic relation between alpha oscillations and firing rates was observed where firing rates were found to be lowest and highest at the peak and trough of alpha oscillations, respectively.

## INTRODUCTION

The brain is a highly adaptable learning machine, and at least partially learns via reinforcement learning mechanisms. As a main function of the brain is to move the individual through their environment in a manner that maximizes reward and minimizes punishment, we hypothesized that one would see neural dynamics reminiscent of various aspects of a reinforcement learning machine (Tarigoppula et al., 2018), and we expected to see this at every level of the neural representation from mesoscopic/macroscopic EEG (i.e., electroencephalography) down to single units (Marsh et al., 2015; McNiel et al., 2016; An et al., 2018), and conducted this work to explore this hypothesis.

Recently, cross-frequency coupling (CFC) has been utilized to measure statistical correlations between different frequency bands of macroscopic (slow frequency oscillations) and/or mesoscopic (high frequency oscillations) local field potential (LFP) oscillations (Soltesz and Deschenes, 1993; Bragin et al., 1995). CFC has been considered an important measure of cognitive information processing in different brain regions (Canolty and Knight, 2010). Phase-to-phase coupling (Palva et al., 2005; Belluscio et al., 2012), phase-to-amplitude coupling (Mormann et al., 2005; Canolty et al., 2006; Tort et al., 2008; Colgin et al., 2009), and amplitude-to-amplitude coupling (Bruns et al., 2000) are different methods in which CFC can be analyzed (Jensen and Colgin, 2007; Canolty and Knight, 2010). In particular, phase-to-amplitude coupling (PAC), between the amplitude of high frequency oscillations and the phase of low frequency oscillations has been observed in cognitive tasks in both human (Canolty et al., 2006; Axmacher et al., 2010) and animal models (Buzsaki et al., 2003; Lakatos et al., 2005; Tort et al., 2008). In addition, the correlation between microscopic (spike trains) and macroscopic and/or mesoscopic (LFPs) scales has been considered to play an important functional role in neural processing (Fries et al., 2001; Jarvis and Mitra, 2001). Several studies have employed spike-field coherence (SFC), the phase dependency between spikes and LFPs, to investigate the communication and synchronization within neuronal groups of the same or different cortical regions (Womelsdorf et al., 2006; Witham et al., 2007; Fries et al., 2008; Pesaran et al., 2008; Gregoriou et al., 2009; Jutras et al., 2009; Chalk et al., 2010).

For many years, alpha (8-14 Hz) oscillations were thought to serve a non-functional purpose, where the power in this band reflected the opened or closed state of the eyes (Adrian, 1934). However, several recent studies have shown the functional role of alpha oscillations and their importance in cognitive processing. Specifically, alpha oscillations reflect inhibitory activity in visual and auditory attention (Foxe et al., 1998; Thut et al., 2006; Rihs et al., 2009; Kerlin et al., 2010), perception (VanRullen and Koch, 2003), and working memory (Jensen et al., 2002; Sauseng et al., 2005; Jensen and Mazaheri, 2010) tasks. However, it is less clear how reward expectation affects these oscillations in the primary motor cortex (M1).

In order to better understand reward-related effects on M1, we conducted the present study where we simultaneously recorded neural spiking activity (single-and multi-units) and LFPs from contra/ipsilateral M1 in non-human primates (NHPs) while they performed cued trial value center-out reaching tasks and grip force tasks. We have previously shown that reward modulates single unit activity, population firing rates, and LFP power in M1 while NHPs either performed or observed a single target center-out reaching task (Marsh et al., 2015), work that has since been corroborated and extend by others (Ramkumar et al., 2016; Ramakrishnan et al., 2017). To further understand the impact of reward expectation on other aspects of M1 neural activity, we studied power spectral density, phase-to-amplitude comodulation, spike-field coherence, and phase correlation with spike firing of neural signals.

## MATERIALS AND METHODS

### Surgery

A rhesus macaque (NHP P (female): Macaca mulatta) and three bonnet macaques (NHPs A (male), S (male), and Z (female): Macaca radiata) were implanted with 96-channel microelectrode Utah arrays (10 × 10 array consisting of 1.5 mm length electrodes spaced 400 micrometer, Blackrock Microsystems, LLC.) in the M1 region associated with their right hand and forearm. NHPs A, S, and P were implanted in the contralateral M1 with respect to the right arm, which all NHPs used to perform the manual tasks. NHP Z was previously implanted twice in the contralateral M1, therefore the array was implanted in ipsilateral M1 for this study. All studies and procedures were approved by the Institutional Animal Care and Use Committee (IACUC) at the State University of New York (SUNY) Downstate Medical Center and complied with the National Institutes of Health Guide for the Care and Use of Laboratory Animals guidelines. The surgical procedures used in the experiment were the same as those as described in our previous work (Chhatbar et al., 2010; Marsh et al., 2015). In brief, veterinary staffs from the SUNY Downstate Division of Comparative Medicine performed general anesthesia and animal preparation. Aseptic conditions were maintained during the course of surgery. Anesthesia was induced with ketamine and maintained using isoflurane and fentanyl. To prevent inflammation, dexamethasone was used during the surgical procedure.

The first surgery for each animal was the implantation of a back post, a titanium post (Crist Instrument Co., Inc.) implanted onto the caudal aspect of the NHP’s cranium to attach to the primate chair during training and recording. The post was placed on the caudal aspect of the skull, holes drilled and tapped (Synthes 2.0 mm drill bit, Synthes 2.0 mm tap) and affixed with 6mm or 8mm screws (Synthes, 2.7mm diameter, titanium) depending on the thickness of the NHP’s skull.

For microelectrode implantation the animal was prepared for surgery in the same way. A rostrally placed front post (titanium, Crist Instrument Co., Inc.) was affixed similarly to the back post, to serve as a platform for the electrode connectors. An approximately 2 cm by 3 cm craniotomy window was then created using a dremel tool with conical tip (Dremel Multipro), over the cortical areas of interest. The dura mater was reflected, and the target locations were identified visually with cortical landmarks. To confirm the location of S1, an electrode (Michigan probe, 4-shank 32 channel silicone array, NeuroNexus Technologies, Inc.) was lowered stereotactically into the cortex. A lab member then stimulated the animal by tapping the contralateral hand and arm, with the electrode output represented audibly through loudspeakers routed through a TDT recording system (Tucker Davis Technologies, Inc.). A 96-channel microelectrode array (Blackrock Utah array, platinum/iridium, 1.5 mm in length) was placed in the determined S1 location, and a pneumatic piston (pneumatic control box, Cyberkinetics Neurotechnology Systems, Inc.) used to secure the array into place in the cortex. The M1 implanted location was immediately across the central sulcus, and put into place in the same manner.

Electrode wires were gathered together and routed to one corner of the craniotomy, and then along the front post to the electrode connectors, which were placed into a plastic frame attached to the front post. Once all the connectors were affixed the dura was sutured back into place, and the bone fragment from the craniotomy window placed above. The bone was affixed with a titanium mesh and bone screws (1.9mm diameter, 4mm in length titanium self-tapping screws, Bioplate Inc.) on the skull and on the bone fragment. Dental acrylic (Palacos, Zimmer Biomet) was used to attach the electrode wires to the skull, and additionally to create a protective layer between the craniotomy and the front post; so, the NHP could not interfere with the wires. NHPs were given a six week rest period following back post implantation to allow for osseointegration before resuming training and recording, and two weeks following microelectrode implantation to allow for the site to heal and cortical inflammation to reduce.

### Cued Center-Out Reaching Task

Two NHPs (A and Z) were trained to perform a center-out delayed hold reaching task with their right arm using a two-link robotic exoskeleton (KINARM, BKIN Technologies Ltd.) for the manual task, as shown in Figure 1a. They were also trained to observe a feedback cursor moving automatically with a constant speed toward the target without performing physical effort during the observational task, as shown in Figure 1b. During the manual task, the NHPs would have to hold their hand on a central target for 325 ms, following a color cue period of 100 – 300ms dependent on the NHPs temperament. During the color cue period the peripheral targets color and the hold target’s color indicated the trials value that is rewarding or non-rewarding. The NHP had to wait another 325-400ms until the go cue, which was the disappearance of the hold target. For a successful trial, the NHP had to reach and hold on the peripheral target for 325 ms. Every successful reach in rewarding trials resulted in a juice reward to the NHP, whereas reward was withheld on nonrewarding successful trials. If a nonrewarding trial was unsuccessful, it was repeated to encourage NHPs to make successful movements. During observational tasks, NHPs observed passively while a feedback cursor moved automatically from center to peripheral target at a constant speed (approximately 1 cm per second). Visual color cues were similar to the manual task, and informed the NHP about the rewarding or nonrewarding trial value if successful. During manual and observational tasks, rewarding and nonrewarding trials were presented in a random order. There was one exception for NHP A during observational tasks where trials followed a set structure or rewarding followed by nonrewarding and repeating.

**Figure 1.**
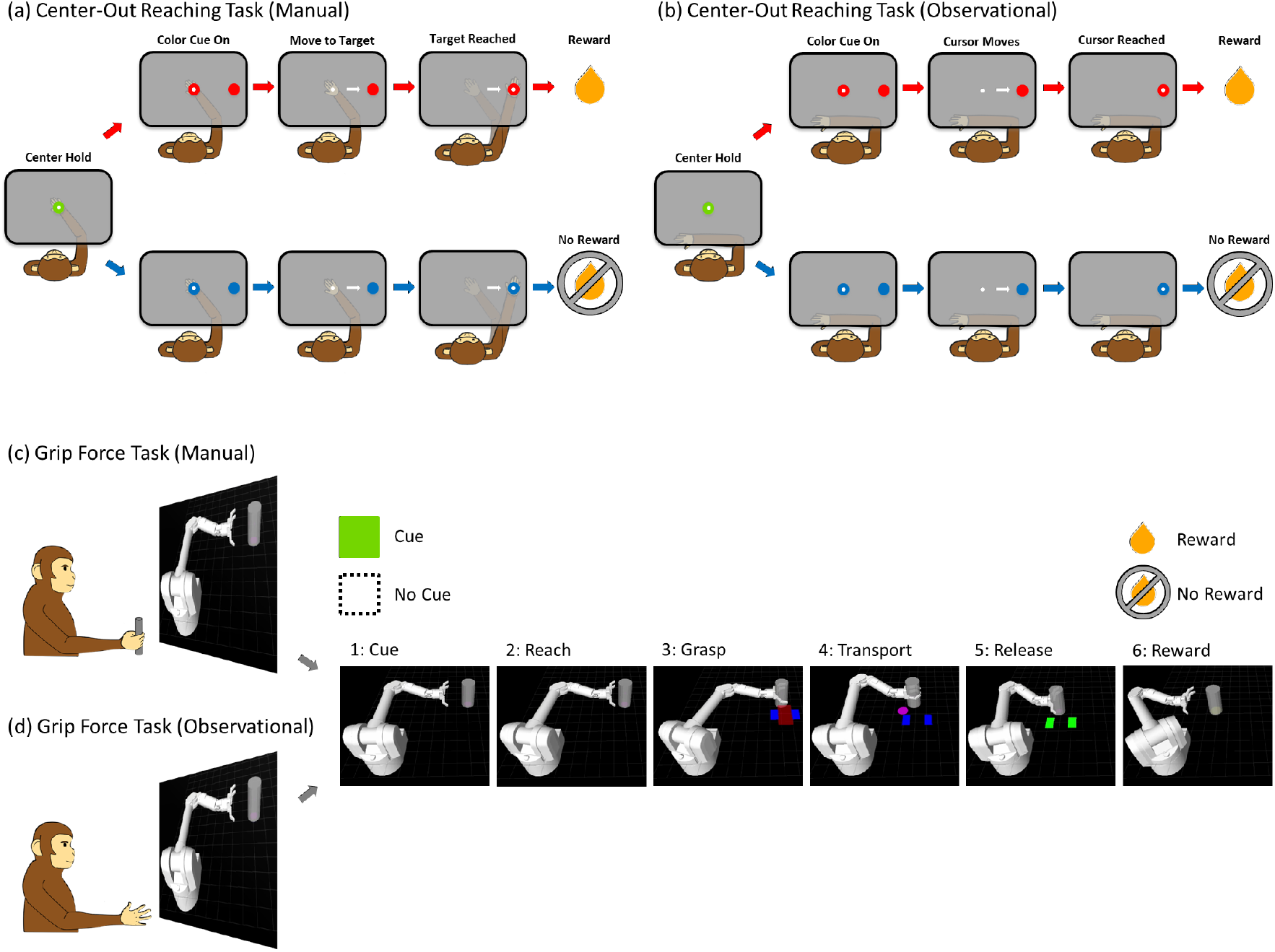
Center-out reaching tasks and grip force tasks. Schematic of (a) manual and (b) observational task during the cued center-out reaching tasks as well as for cued manual (c) and observational (d) grip force tasks.

### Cued Grip Force Task

Two NHPs (S and P) were trained to perform cued grip force tasks sitting comfortably in a primate-training chair (BKIN Technologies Ltd.) either manually, Figure 1c, or observationally, Figure 1d. The task consisted of a virtual robotic arm, modeled as a Barrett WAM Arm and Hand (Barrett Technology) in Gazebo on ROS (Robot Operating System) that reached toward a virtual cylindrical object. When the robotic arm reached the object, the NHP was required to apply and maintain a visually indicated amount of force on a custom-made manual gripper (force transducer) which was composed of a metal bar, a cylindrical plastic frame, and a force transducer (FC2231-0000-0100-L, Measurement Specialties, Inc.) to measure grip force while the cylinder was transported to the target position by the WAM simulation. Once at the target position, the NHP would release the manual gripper, and the robotic hand would release the object, and the arm moved back to the neutral starting position. The target force was indicated with a pair of blue rectangles in the virtual environment, where the width of the rectangles represented the upper and lower bounds of the required grip force. The actual magnitude of force applied by the NHP was shown as a red rectangle that expanded as the force increased. The correct amount of grasping force was maintained by keeping the red rectangle within the upper and lower bounds of the blue rectangle. For a trial to be considered successful two conditions had to be met. The first was the application of appropriate force during the transport of the object to the target location, and the second condition was the release at the end of object’s transfer. Based on whether a trial was cued rewarding or nonrewarding, the NHP respectively received or did not receive a juice reward at the end of a successful trial. The visual color cue was shown at the beginning of each trial and remained visible throughout the trial. This trial progression was divided into six timeframes for analysis: (i) Cue, when the cue was presented; (ii) Reach, when the virtual arm approached the target; (iii) Grasp, when the NHP applied force to the manual gripper so the virtual hand grasped the object; (iv) Transport, where the object automatically moved to the target location while the NHP maintained grip force; (v) Release, when the gripper was released and the virtual hand released the object; and (vi) Reward or Nonreward, when the NHP received a juice reward for successful completion of a rewarding trial, did not receive reward for completion of a nonrewarding trial, or did not receive reward for the unsuccessful completion of either trial type.

The structure of rewarding and nonrewarding trials was completely predictable in some recording blocks, where trials alternated between the two. The remainder of the blocks were partially predictable, where 50%, 75%, or 90% of the trials were rewarding, and the trial type was selected pseudorandomly with this bias. In the observational task, the mechanics of the task were the same, but the NHP had to passively observe the robot performing an automatic grasp and transport of the cylinder instead of manually applying force.

### Neural Recording

Spike trains and LFPs were recorded simultaneously from M1 cortices using a Multichannel Acquisition Processor recording system (MAP, Plexon Inc.) while the NHPs performed center-out reaching tasks and grip force tasks. The recorded neural signals were bandpass-filtered from 170 Hz to 8 kHz for spike trains and from 0.7 to 300 Hz for LFPs using a linear finite impulse response (FIR) filter, and sampled at 40 kHz for spikes and 2 kHz for LFPs using a MAP system. Offline spike sorting was performed using a custom-made sorting template in the commercial software, Offline Sorter (Plexon Inc.). We analyzed single/multiunit activity and LFPs recorded from the contralateral M1 area of NHPs A, S and P as well as the ipsilateral M1 area of NHP Z.

### Analysis of Power Spectral Density

To probe whether reward expectation modulates the power of neural oscillations, power spectral density (PSD) of the LFPs was estimated using the Welch periodogram method with 75% overlapping Hamming windows (Welch, 1967). LFPs recorded from 32 channels were preprocessed by first removing the line noise (60 Hz) from every LFP channel using a second-order Butterworth notch filter, following which each channel was z-scored. Then, LFP channels with signal-to-noise ratio (SNR) below 5 were excluded from further analysis. SNR was calculated as the ratio of peak-to-peak amplitude (*A_Pk-Pk_*) of the averaged LFP (*LFP_Avg_*) and twice the standard deviation (SD) of the residual signal (given by the difference of LFP on *i^th^* channel and *LFP_Avg_*) as in Equation (1) where *i ∈ [1,32]* (Koralek et al., 2013)

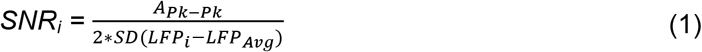

This SNR calculation was performed separately for each NHP dataset. Based on SNR criterion, the number (average ± standard deviation) of LFP channels used for manual task datasets was 27.5 ± 1.0, 26.0 ± 2.7, 30.7 ± 0.6, and 24.0 ± 0.0 in NHPs A, Z, S and P, respectively. For observational task datasets, 24.0 ± 2.0, 24.3 ± 3.5, 30.0 ± 0.0, and 27.0 ± 5.8 of LFP channels were used for NHPs A, Z, S and P, respectively. Selected LFP channels were averaged and used for PSD estimation. For comparison between rewarding and nonrewarding trials, each PSD was then normalized by dividing the power at each frequency by the average of all power from 0.5 to 100 Hz (Colgin et al., 2009). The trial-averaged PSD in the alpha band (8 to 14 Hz) was employed to compare rewarding to nonrewarding trials while NHPs performed all tasks.

### Analysis of Cross-Frequency Coupling

To quantify the cross-frequency coupling in LFPs and test its relationship with reward expectation, the phase-amplitude coupling (PAC) method (Kullback-Leibler based modulation index (MI)) was used as proposed by Tort et al. (Tort et al., 2008). PAC analysis can be described briefly in the following manner (Tort et al., 2008). PAC appears when the amplitude of fast frequency oscillations is modulated by the phase of slow frequency oscillations (Lakatos et al., 2005; Canolty et al., 2006; Jensen and Colgin, 2007; Tort et al., 2008). First, the averaged LFP was bandpass filtered at a low frequency band (8 Hz to 20 Hz) for phase and a high frequency band (25 Hz to 100 Hz) for amplitude. Second, the phase of the low frequency band and amplitude of the high frequency band were extracted from the above filtered LFP by applying Hilbert transform. Third, the strength of amplitude comodulation by phase was computed as the MI measure at each phase-frequency and amplitude-frequency pair. The MI was calculated in steps of 0.5 Hz for phase-frequency and 5 Hz for amplitude-frequency. Using this PAC method, we computed the strength of phase-to-amplitude comodulation in rewarding and nonrewarding trials for center-out reaching tasks and grip force tasks, and compared them for statistically significant differences.

### Analysis of Spike-Field Coherence

To examine whether reward expectation modulates the coherence between spike trains and LFPs, we conducted spike-field coherence (SFC) analysis. SFC is commonly used to compute phase synchronization between point process data (e.g., spike trains) and continuous data (e.g., LFPs). To rule out the possibility that spikes might have contaminated LFPs (Waldert et al., 2013), all LFPs from selected channels (higher SNR channels) were averaged, then the power spectra of the binned point processes (single unit activity, binned at 1 ms) and channel-averaged LFP were computed using the multitaper estimation method. For this purpose, we used a Chronux built-in function, *coherencycpb(),* with 9 tapers and a time-bandwidth (TW) value equal to 5. The coherence between spikes and LFPs was computed using the formula as shown in Equation (2) (Fries et al., 2001; Jarvis and Mitra, 2001)

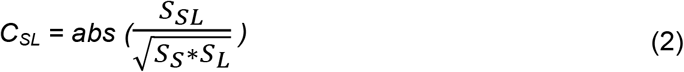

Here, *S_SL_* indicates the cross-spectrum between spikes and LFPs, and *S_S_* and *S_L_* indicate the spectra of spikes and LFPs respectively. An SFC value (*C_SL_*) of zero indicates the absence of phase synchronization between spikes and LFPs, and one indicates the perfect synchronization between them. The trial-averaged SFC was analyzed over the frequency band (0.5 to 100 Hz) and the trial-averaged SFC at the alpha frequency band was used to compare rewarding trials to nonrewarding trials while NHPs performed the aforementioned tasks.

### Relation between Alpha Cycle and Neural Spiking

The interaction between the alpha-band cycle and neural spiking activity was measured as follows (Haegens et al., 2011): first, we bandpass filtered LFP oscillations from 8 Hz to 14 Hz; second, the phase of the alpha-band LFP was obtained by applying the Hilbert transform; third, we divided the alpha-band cycle into six equally sized phase bins of 60 degrees each; fourth, each unit’s firing rate (FR) was normalized across the six phase bins using the min-max scaling method, obtained by *Normalized FR = [FR – min(FR)] / [max(FR) – min(FR)];* last, the average firing rate in each phase bin was computed over a population of units and represented with respect to the corresponding phase of alpha oscillations.

### Statistical Analysis

All statistical analyses were performed using MATLAB R2017b (MathWorks Inc.). The nonparametric Wilcoxon signed rank test (i.e., *signrank()*) was used to evaluate significant differences of PSD, PAC, SFC, and spike firing rates between rewarding and nonrewarding trials for all tasks. To test the statistical significance of changes in firing rate with respect to the alpha-band phase bins, we performed one-way ANOVA with Bonferroni post-hoc multiple comparison (significance level: α = 0.05) for rewarding and nonrewarding trials individually. The F-statistic value (from one-way ANOVA) and p-value computed between the phase bins of low and high firing rates (from two-sample *t*-test) were observed for different tasks and trial types.

## RESULTS

In order to investigate how cued reward expectation influences neural activity in M1 cortex, we performed PSD and PAC analyses with LFPs, computed spike firing rate, and quantified SFC on neural spikes and LFP data obtained from contralateral (NHPs A, S and P) and ipsilateral (NHP Z) M1 across center-out reaching tasks and grip force tasks. In addition, we investigated how alpha band oscillations correlate with spike firing rate. As mentioned in the methods, task structure could either be completely predictable or random with a bias. For NHP A’s observational task, rewarding and nonrewarding trials were presented as a sequence alternating between the two while all the other tasks for all NHPs were random with a bias which varied based on the percentage of reward bias.

### Reward Expectation Modulates Alpha Power

For PSD analysis, we used a post-cue-onset period of 800 ms for all tasks. Figure 2 displays the normalized PSD plots (left column in each subplot) and bar plots for the alpha band (right column in each subplot). Shown are significant differences of alpha (8 to 14 Hz) LFPs for rewarding (red) and nonrewarding (blue) trials across manual (upper row in each subplot) and observational (lower row in each subplot) tasks during both center-out reaching tasks (NHPs A and Z as shown in Fig. 2a and 2b) and grip force tasks (NHPs S and P as shown in Fig. 2c and 2d).

**Figure 2.**
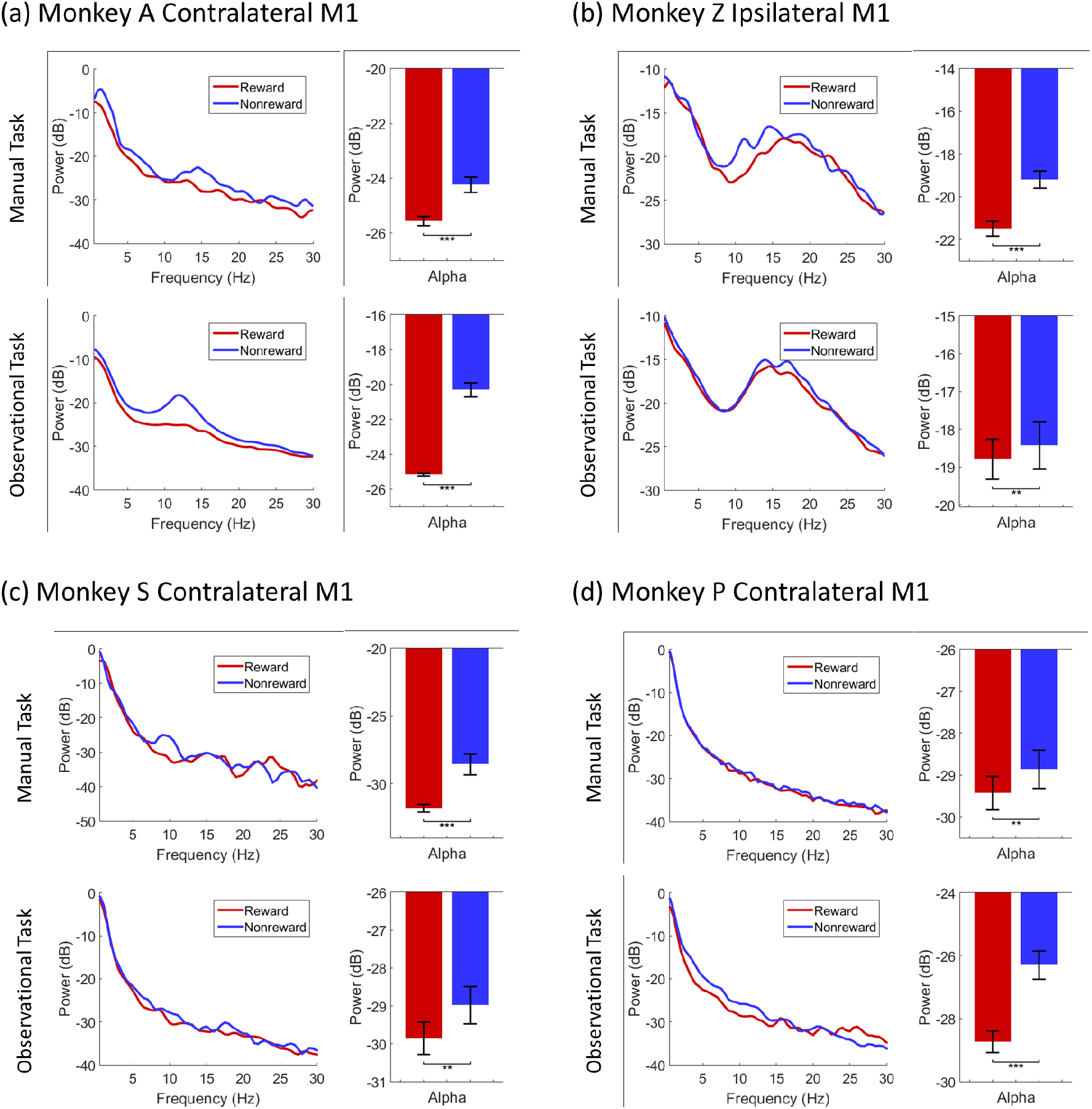
Power spectral density (PSD) plots (left column in each subplot) of LFPs for rewarding (red) and nonrewarding (blue) trials across manual (upper row in each subplot) and observational (lower row in each subplot) tasks during center-out reaching (a and b) tasks and grip force (c and d) tasks. Bar graphs show significant differences of alpha (8-14 Hz, right column in each subplot) band (**** denotes p < 0.0001; *** denotes p < 0.001; ** denotes p < 0.01; * denotes p < 0.05; Wilcoxon signed rank test). Error bars in the bar plots represent SEM.

During the manual reaching task from NHP A’s contralateral M1 (Fig. 2a upper row), the PSD in the alpha band was significantly higher during nonrewarding trials than rewarding trials (Wilcoxon signed rank test, *p* < 0.001). Furthermore, for the observational task (Fig. 2a lower row), we found similar patterns of neural activation. Similar to the manual task, alpha band PSD was significantly higher during nonrewarding than rewarding trials for the observational task (Wilcoxon signed rank test, *p* < 0.001). During the manual-reaching task from NHP Z’s ipsilateral M1 (Fig. 2b upper row), again the alpha power was significantly higher during nonrewarding than rewarding trials (Wilcoxon signed rank test, *p* < 0.001). During the observational task (Fig. 2b lower row), alpha PSD showed a significant increase during nonrewarding trials as well (Wilcoxon signed rank test, *p* < 0.01).

Data from the manual grip-force task from NHP S’s contralateral M1 (Fig. 2c upper row) showed that alpha PSD was significantly higher for nonrewarding than rewarding trials (Wilcoxon signed rank test, *p* < 0.001). Similarly, for the observational task (Fig. 2c lower row), alpha PSD was significantly higher for nonrewarding than rewarding trials (Wilcoxon signed rank test, *p* < 0.01). During both manual (Fig. 2d upper row) and observational (Fig. 2d lower row) grip-force tasks from NHP P’s contralateral M1, alpha PSD was significantly higher for nonrewarding than rewarding trials (Wilcoxon signed rank test, *p* < 0.01 for manual and *p* < 0.001 for observational). The overall PSD results in Figure 2 indicate that the alpha power of LFP oscillations were modulated by cued reward expectation. It also indicates that alpha power was significantly increased for nonrewarding trials as compared to rewarding trials for both manual and observational variations of arm reaching tasks and hand grasping tasks.

### Reward Expectation Modulates Alpha-Gamma Comodulation

To determine whether the phase of low frequency oscillations was related to the amplitude of high frequency oscillations in M1 during our reward cued experiments (see Fig. 1), we computed phase-to-amplitude comodulation during a post-cue-onset period of 800 ms both for rewarding and nonrewarding trials across all tasks. Figure 3 displays phase-to-amplitude comodulogram plots for rewarding trials (left column in each subplot) and nonrewarding trials (middle column in each subplot) across manual (upper row in each subplot) and observational (lower row in each subplot) tasks for contralateral (Fig. 3a, 3c and 3d) and ipsilateral (Fig. 3b) M1 cortices. In addition, the bar plot distributions (right column in each subplot) showed significant differences of alpha-gamma comodulation index values for rewarding (red) and nonrewarding (blue) trials. During the manual reaching task from NHP A’s contralateral M1 (Fig. 3a upper row), alpha-gamma PAC was significantly greater for nonrewarding trials than rewarding trials (Wilcoxon signed rank test, *p* < 0.0001) when the phase is in the alpha frequency at approximately 10 Hz<. For the observational task (Fig. 3a lower row), alpha-gamma PAC was also significantly higher during nonrewarding trials than during rewarding trials (Wilcoxon signed rank test, *p* < 0.0001) at alpha phase within the frequency band of 10-14 Hz. Similar trends were observed for NHP Z’s ipsilateral M1 (Fig. 3b), where alpha-gamma comodulation at phase frequencies of 8-10 Hz for both manual and observational tasks during nonrewarding trials were significantly greater than rewarding trials (Wilcoxon signed rank test, *p* < 0.0001, for both manual and observational tasks). The results for the center-out reaching tasks indicate that the strength of alpha-gamma frequency PAC analyzed with the averaged LFP oscillations recorded from contralateral and ipsilateral M1 were modulated by reward expectation in both the presence (for manual) and absence (for observational) of arm reaching movements.

**Figure 3.**
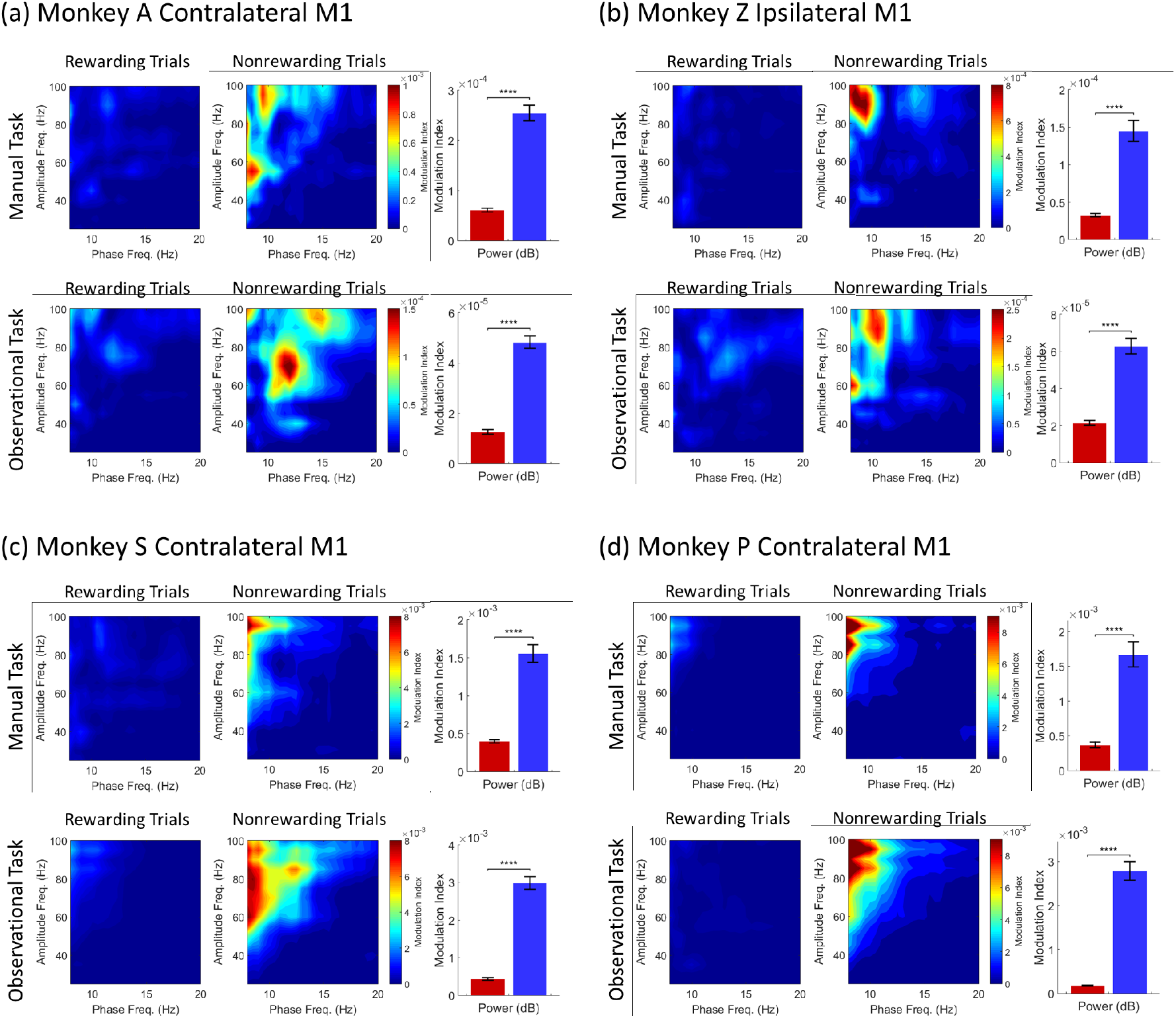
Comodulograms showing the modulation index (MI) for rewarding (left column in each subplot) and nonrewarding trials (middle column in each subplot) across manual (upper row in each subplot) and observational tasks (lower row in each subplot) for contralateral (a, c and d) and ipsilateral (b) M1 cortices. Bar graphs (right column in each subplot) show significant differences of alpha-gamma comodulation index values for rewarding (red) and nonrewarding (blue) trials. (**** denotes p < 0.0001; *** denotes p < 0.001; ** denotes p < 0.01; * denotes p < 0.05; Wilcoxon signed rank test). Error bars in the bar plots represent SEM. (a and b) are for the center-out reaching tasks, and (c and d) are for the grip force tasks.

Qualitatively similar PAC results were observed in the grip force tasks. In Figure 3c, alpha-gamma comodulations for both the manual (upper row) and observational (lower row) grip-force tasks from NHP S’s contralateral M1 were significantly higher during nonrewarding trials than rewarding trials (Wilcoxon signed rank test, *p* < 0.0001, for both manual and observational tasks). In Figure 3d, the PAC results of NHP P also showed significantly higher alpha-gamma comodulation for nonrewarding trials than rewarding trials for both the manual (upper row) and observational (lower row) tasks (Wilcoxon signed rank test, *p* < 0.0001, for both manual and observational tasks). Similar to PAC results for the center-out reaching tasks, our results demonstrate that the strength of alpha-gamma comodulation was influenced by reward expectation during both manual and observational grip force tasks.

### Reward Expectation Modulates Spike-Field Coherence

To investigate whether alpha oscillations are related to phase synchronization between spikes and LFPs, reward-related changes in the alpha band (8 to 14 Hz) SFC were estimated during the center-out reaching tasks and grip force tasks. Figure 4 shows SFC plots (upper row in each subplot) for sample units (with specific unit number) during a postcue (after cue) period of 800 ms for rewarding (red) and nonrewarding (blue) trials for contralateral (NHPs A, S and P) and ipsilateral (NHP Z) M1 cortices across all tasks. Additionally, Figure 4 displays the population (lower row in each subplot) of significantly different units for SFC values in the alpha band during rewarding (red) and nonrewarding (blue) trials, or neither (gray) (Wilcoxon signed rank test, p < 0.05). From left to right in the bar charts (lower row in each subplot), each column represents the precue (before cue, 500 ms), postcue (after cue, 800 ms), prereward (before reward, 500 ms), and postreward (after reward, 500 ms) time windows. In NHP A’s contralateral M1 during manual center-out reaching tasks (left column in Fig. 4a), 37.8% (133 of 352 for precue), 58.0% (204 of 352 for postcue), 51.7% (182 of 352 for prereward), and 39.2% (138 of 352 for postreward) of M1 units had significantly higher trial-averaged SFC during nonrewarding trials than rewarding trials. For observational center-out “reaching” tasks (right column in Fig. 4a), the percentages are 22.0% (81 of 367 for precue), 71.9% (264 of 367 for postcue), 47.7% (175 of 367 for prereward), and 38.4% (141 of 367 for postreward) of M1 units. Similar results were seen in NHP Z’s ipsilateral M1 during the center-out reaching tasks. The detailed breakdown of the results can be seen in Figure 4b (left column for manual task and right column for observational task).

**Figure 4.**
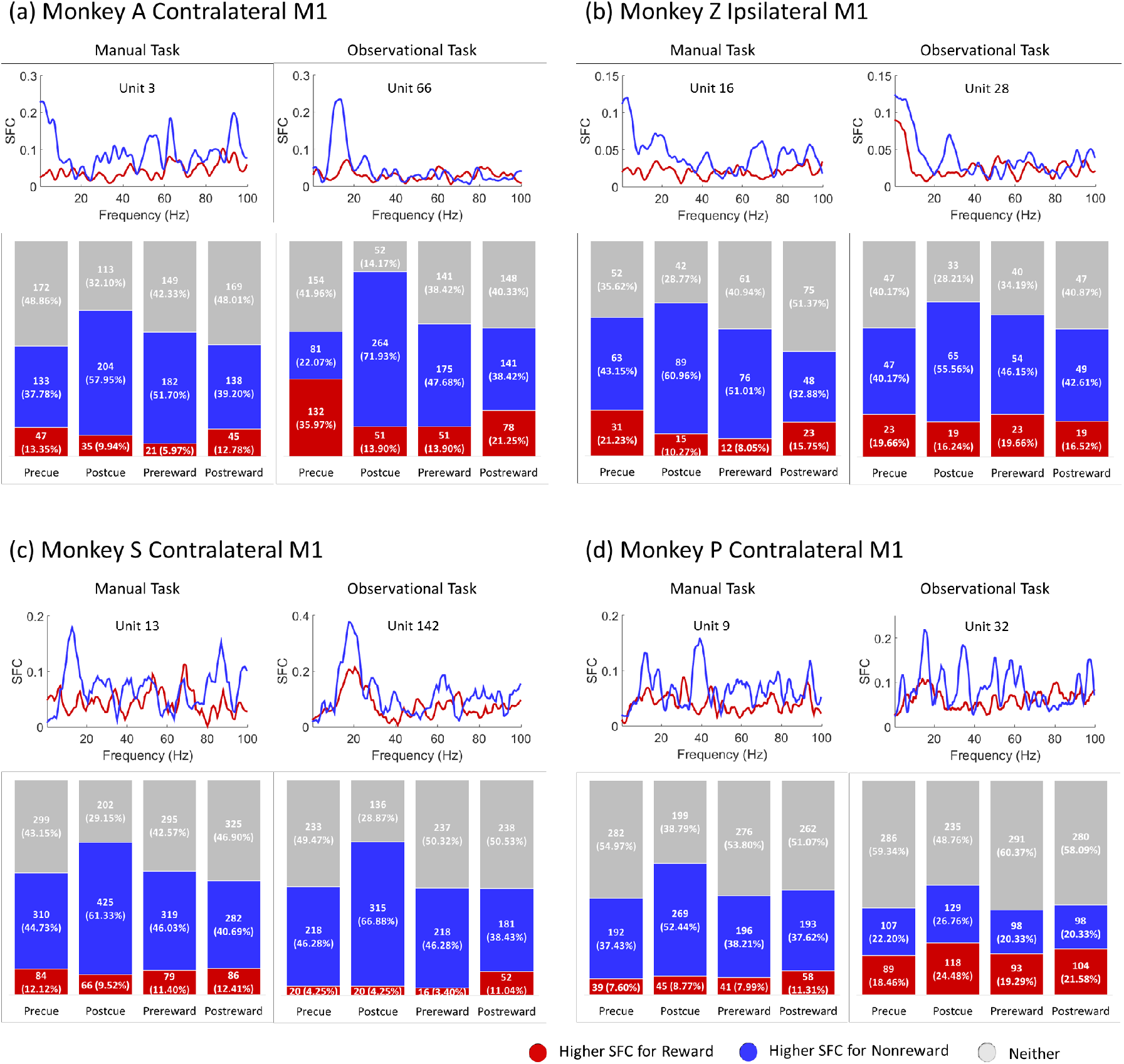
Spike-field coherence (SFC) plots (upper row in each subplot) for sample units for rewarding (red) and nonrewarding (blue) trials across manual (left column in each subplot) and observational (right column in each subplot) tasks for contralateral (a, c, and d) and ipsilateral (b) M1 cortices. Bar charts (lower row in each subplot) represent the population of significantly different M1 units for SFC values in alpha (8 to 14 Hz) band during rewarding (red) and nonrewarding (blue) trials, and those with no significant difference (gray) (Wilcoxon signed rank test, p < 0.05). Each column in the bar chart represents precue (before cue, 500 ms), postcue (after cue, 800 ms), prereward (before reward, 500 ms), and postreward (after reward, 500 ms) periods. (a and b) are for the center-out reaching tasks, and (c and d) are for the grip force tasks.

As with NHPs A and Z, NHP S also showed significantly higher SFC percentages in nonrewarding trials for both manual and observational grip force tasks shown in Figure 4c. This pattern remains consistent in NHP P for manual tasks (left column in Fig. 4d), but with less significance for observational tasks (right column in Fig. 4d). Furthermore, the postreward (after reward delivery) period in observational tasks for NHP P showed a slightly higher percentage of SFC for rewarding than nonrewarding trials. Overall, the results in Figure 4 indicate that M1 units are significantly modulated by reward expectation for alpha-band SFC during all reaching tasks and grasping tasks.

### Reward Expectation Modulates Spike Firing Rate

To examine the reward expectation modulation of neural spiking in M1, we computed the firing rate of M1 units during the center-out reaching tasks and grip force tasks. For each M1 unit, firing rate was computed using 50 ms bins during the following periods, precue (500 ms), postcue (800 ms), prereward (500 ms), and postreward (500 ms). Figure 5 displays the total percentage of M1 units that have significantly higher average firing rates for rewarding (red) trials and nonrewarding (blue) trials (Wilcoxon signed rank test, *p* < 0.05). In NHP A’s contralateral M1 (left column in Fig. 5a), 11.1% (39 of 352 for precue), 36.1% (127 of 352 for postcue), 26.7% (94 of 352 for prereward) of M1 units for the manual task had significantly higher firing rates during rewarding trials than nonrewarding trials; whereas, 19.6% (69 of 352 for postreward) of M1 units had significantly higher firing rates for nonrewarding trials than rewarding trials. For the observational task (right column in Fig. 5a), 48.5% for precue and 52.3% for postcue of M1 units had significantly higher firing rates for rewarding trials than nonrewarding trials; whereas, 41.7% for prereward and 51.2% for postreward of M1 units had significantly higher firing rates for nonrewarding trials as compared to rewarding trials. The observation task performed by NHP A was a fully predictable sequence of rewarding trials followed by nonrewarding trials, which explains the precue activity pattern (see (Tarigoppula et al., 2018)). In NHP Z’s ipsilateral M1, during the manual task (left column in Fig. 5b), 19.2% for precue, 37.7% for postcue, and 33.6% for prereward had significantly higher spiking rates for rewarding trials than nonrewarding trials, but 31.5% for postreward period had higher firing rates for nonrewarding trials than rewarding trials. Similar to the manual task, NHP Z showed a similar trend of population firing rates as for the observational task. The detailed population firing rates are shown in Figure 5b.

**Figure 5.**
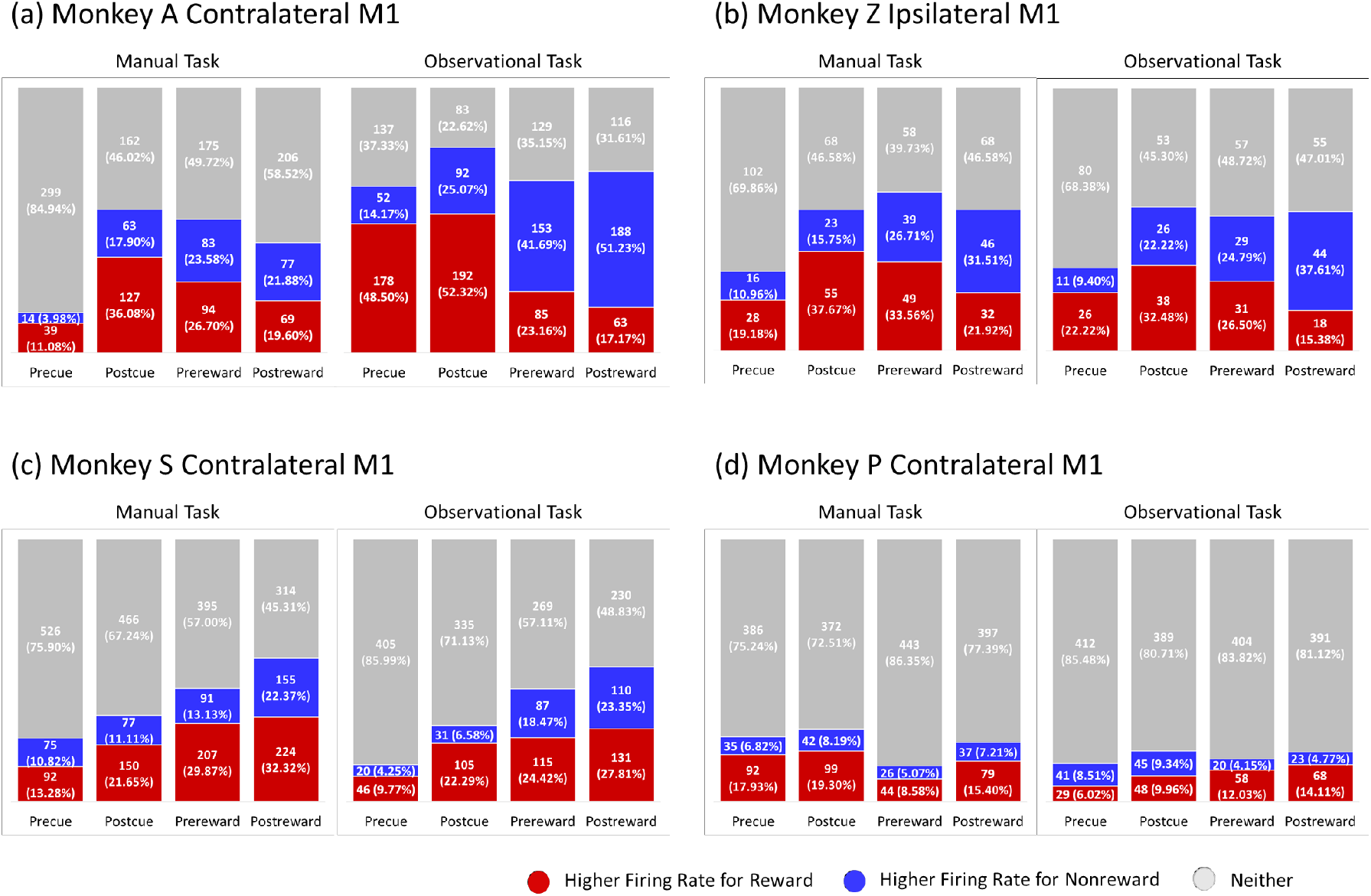
Total population of M1 units that had significantly higher firing rates for rewarding (red), nonrewarding (blue) trials, or neither (gray) across manual (left column) and observational (right column) tasks for contralateral (a, c, and d) and ipsilateral (b) M1 cortices (Wilcoxon signed rank test, p < 0.05). Each column in the bar chart represents precue (500 ms), postcue (800 ms), prereward (500 ms), and postreward (500 ms) periods. (a and b) are for the center-out reaching tasks, and (c and d) are for the grip force tasks.

During grip force tasks, in NHP S’s contralateral M1 results (Fig. 5c), rewarding (red) trials as a whole always showed significantly higher firing rates than nonrewarding (blue) trials for both the manual task (left column) and the observational task (right column). In NHP P’s contralateral M1 (Fig. 5d), during the manual task (left column), firing rates for all periods were significantly higher for rewarding (red) trials than nonrewarding (blue) trials (Wilcoxon signed rank, *p* < 0.05). For the observational task (right column) from NHP P’s contralateral M1 units, all periods except the precue period had higher firing rates for rewarding trials than nonrewarding trials. The number of significant M1 units of NHP P’s neural spiking rate during observational tasks was lower compared with significance of manual tasks. Overall, the results show that neural rates are significantly higher in M1 when reward is expected.

### Alpha Phase of LFP Relates to Neural Spiking Activity

Previous work has shown a significant relationship between alpha power and neural firing rate such that firing rates were highest at the trough and lowest at the peak of the alpha cycle (Haegens et al., 2011). Here we show some support for this observation, and demonstrate that it is strongest during the nonrewarding trials. Figure 6 shows the relationship between the alpha band cycle and spike firing rate for rewarding (left column in each subplot) and nonrewarding (right column in each subplot) trials across manual (upper row in each subplot) and observational tasks (lower row in each subplot) for contralateral (a, c and d) and ipsilateral (b) M1 cortices. In the manual task (top row) for NHP A’s contralateral M1 (Fig. 6a), the firing rate is high around the trough (3π/2) of the alpha band cycle, and low around the peak (π/2) of the cycle for nonrewarding trials (right column; F(5, 660) = 3.95, p < 0.05), whereas this was not observed for rewarding trials (left column; F(5, 744) = 1.36, p > 0.05). Similarly, for the observational task (bottom row in Fig. 6a), firing rate was high around the trough of the cycle, whereas it was low around the peak for nonrewarding trials (right column; F(5, 2040) = 25.79, p < 0.0001). We did not see the same pattern for rewarding trials (left column; F(5, 2040) = 3.7, p < 0.01). In NHP Z’s ipsilateral M1, during the manual task (top row in Fig. 6b), the firing rate was highest around the trough of the alpha cycle for nonrewarding trials (right column; F(5, 534) = 3.35, p < 0.01), but it was highest at the peak and lowest at the trough for rewarding trials (left column; F(5, 564) = 1.17, p > 0.05). During the observational task (bottom row in Fig. 6b), highest firing rates were associated with the trough of the alpha cycle for nonrewarding trials (right column; F(5, 276) = 3.30, p < 0.01) but not for rewarding trials (left column; F(5, 294) = 1.63, p > 0.05).

**Figure 6.**
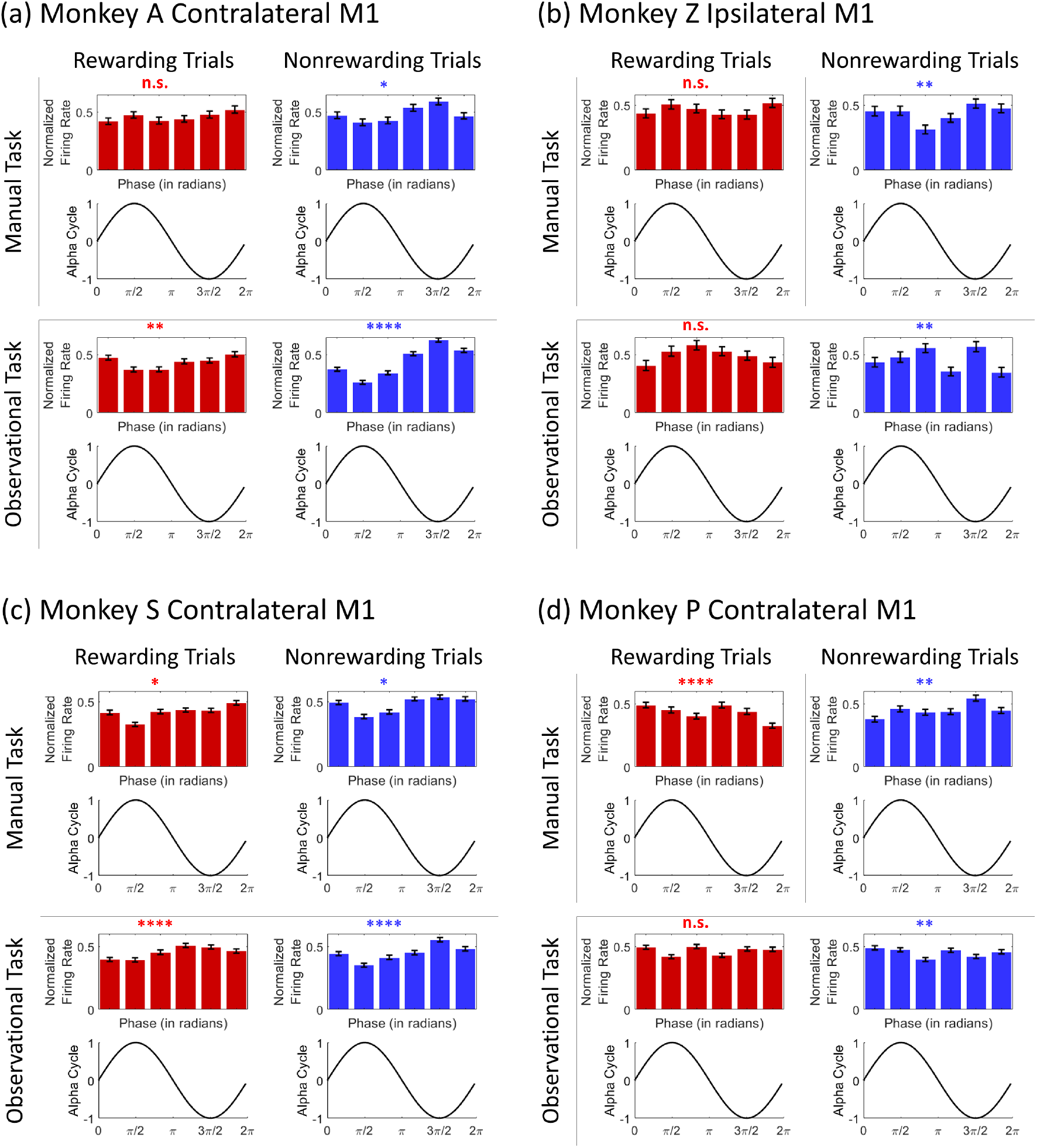
The relationship between neural firing and the alpha band cycle during rewarding (left column in each subplot) and nonrewarding (right column in each subplot) trials across manual (upper row in each subplot) and observational task (lower row in each subplot) for contralateral (a, c and d) and ipsilateral (b) M1 cortices (**** denotes p < 0.0001; *** denotes p < 0.001; ** denotes p < 0.01; * denotes p < 0.05; n.s. denotes no significance; one-way ANOVA). Error bars in the bar plots represent SEM. (a and b) are for the center-out reaching tasks, and (c and d) are for the grip force tasks.

We found similar results in the grip force tasks as seen in center-out reaching tasks. For the manual task (upper row), NHP S’s contralateral M1 (Fig. 6c) showed that the firing rate was high around the trough, and low around the peak of the cycle for nonrewarding trials (right column; F(5, 528) = 2.60, p < 0.05) whereas it was lowest around the peak of the alpha cycle for rewarding trials (left column; F(5, 678) = 2.43, p < 0.05). For the observational task (bottom row in Fig. 6c), firing rate was high at the trough of alpha cycle, and it was low around the peak for nonrewarding trials (right column; F(5, 1890) = 10.68, p < 0.0001). The firing rate was lowest at the peak for rewarding trials (left column; F(5, 2016) = 5.95, p < 0.0001). For NHP P’s contralateral M1 (Fig. 6d), manual task results (upper row) showed that firing rate was highest around the trough but not lowest around the peak for nonrewarding trials (right column; F(5, 1050) = 3.80, p < 0.01). The firing rate does not have the same trend for rewarding trials (left column; F(5, 1020) = 5.04, p < 0.001). For the observational task (bottom row), the highest and lowest firing rates both for rewarding (F(5, 1350) = 1.72, p > 0.05) and nonrewarding trials (F(5,1344) = 3.52, p < 0.01) were not associated with the peak and trough of the alpha cycle.

## DISCUSSION

In order to determine the influence that reward expectation has on M1 in NHPs we recorded neural activity in the form of single and multi-unit activity in conjunction with LFPs from hundreds of chronically implanted electrodes in contra/ipsilateral cortical regions. We utilized multiple sensorimotor tasks where NHPs either made reaching or grasping movements, or observed those movements through cursor motion on a computer screen, or the simulated movement of an anthropomorphic robotic arm reaching and grasping virtual items. We found several clear and reproducible patterns of activity between NHPs, cortical hemispheres and tasks. These patterns of activity included an increase in alpha power in the absence of reward expectation when the trial was cued as being nonrewarding, even though for manual tasks this still required targeted movements by the NHPs. Nonrewarding trials also had much stronger SFC between the neural spiking activity and the averaged LFP activity in the alpha band. There was also a clear increase in phase-amplitude coupling between the alpha phase and gamma amplitude for the nonrewarding trials compared to the rewarding trials.

### Reward-Modulated Changes in Power Spectral Density (PSD)

Previous studies have reported that the power of LFP oscillations is modulated by visual and auditory attention (Foxe et al., 1998; Thut et al., 2006; Rihs et al., 2009; Kerlin et al., 2010) and reward expectancy (van Wingerden et al., 2010; Lansink et al., 2016). Our PSD results showed a consistent reward-related decrease in the mean power of alpha band activity in bilateral M1 cortices during the post-cue-onset period for rewarding trials as compared to nonrewarding trials for both manual and observational tasks, during both reaching tasks and grasping tasks (see Figure 2). Recent evidence suggests that dopamine plays a critical role in the selection of targets for attention as well as in the stabilization of attention against interference in the prefrontal cortex (PFC) of pigeons (Rose et al., 2010). Additionally, injecting a dopamine D1-agonist into the PFC of rats enhanced attentional accuracy, and a D1-antagonist in the same region led to decreased performance (Granon et al., 2000; Chudasama and Robbins, 2004). Furthermore, dopamine depletion has an effect on several neuropsychiatric disorders such as Parkinson’s disease (Cassidy et al., 2002; Sharott et al., 2005; Costa et al., 2006; Mallet et al., 2008; Lemaire et al., 2012), schizophrenia (Abi-Dargham et al., 2000; Arnsten et al., 2017), and attention-deficit/hyperactivity disorder (ADHD) (Jucaite et al., 2005; Silvetti et al., 2013) associated with functional impairments in attention (Posner, 1980; Maunsell, 2004). These studies suggest that changes in dopamine transmission or release could be responsible for changes in attention. Following this, it is expected that subjects may pay more attention to the targets that result in rewards (Dalley et al., 2002; Demiralp et al., 2007; Rose et al., 2010) than targets that do not. In our cued reaching tasks and grasping tasks, subjects may not have paid as much attention during nonrewarding trials as during rewarding trials, potentially due to reduced dopamine release in the absence of reward. We expect to see this difference more clearly in a fully predictable task structure such as NHP A’s observational task. In turn, alpha power was comparatively greater for nonrewarding trials than rewarding trials. Our findings are consistent with previous studies showing that dopamine depletion led to an increased power of LFP oscillations (Cassidy et al., 2002; Sharott et al., 2005; Costa et al., 2006; Kuhn et al., 2008; Mallet et al., 2008; Lemaire et al., 2012). These results provide a clue toward explaining the relationship that exists between dopamine, reward expectation (or attention/motivation), and LFP oscillations in M1 cortex.

### Reward-Modulated Changes in Phase Amplitude Coupling (PAC)

Recently, the phase of low frequency oscillations has been shown to modulate with the amplitude of high frequency oscillations (Canolty et al., 2006; Tort et al., 2008; Canolty and Knight, 2010). Some evidence of phase-to-amplitudes comodulation has come from the many studies conducted across different frequency bands: delta-gamma (Gross et al., 2013; Lopez-Azcarate et al., 2013; Szczepanski et al., 2014), theta-gamma (Bragin et al., 1995; Chrobak and Buzsaki, 1998; Canolty et al., 2006; Tort et al., 2008; Voytek et al., 2010; Lisman and Jensen, 2013; Voloh et al., 2015), alpha-gamma (Osipova et al., 2008; Cohen et al., 2009; Voytek et al., 2010; Spaak et al., 2012; Yanagisawa et al., 2012; van Kerkoerle et al., 2014; Bonnefond and Jensen, 2015; Park et al., 2016; Seymour et al., 2017; Tzvi et al., 2018), and beta-gamma (de Hemptinne et al., 2013; Kim et al., 2015; Swann et al., 2015). In particular, some of these studies investigated alpha-gamma comodulation in the visual cortices (Osipova et al., 2008; Voytek et al., 2010; Spaak et al., 2012; van Kerkoerle et al., 2014; Bonnefond and Jensen, 2015; Seymour et al., 2017), parietal-occipital areas (Tzvi et al., 2018), lingual gyrus (Park et al., 2016), and sensorimotor cortex (Yanagisawa et al., 2012). In our analysis, we found that reward expectation influenced the comodulation between the phase of alpha-band (8–14 Hz) oscillations and the amplitude of gamma-band (30–100 Hz) oscillations during the postcue period in M1 (see Figure 3). We found a higher strength of phase-to-amplitude comodulation in nonrewarding trials during both manual tasks and observational tasks while performing either a reaching (NHPs A and Z) or grasping movement task (NHPs S and P). In addition, in our work, an increase in alpha power had the tendency to lead to stronger alpha-gamma comodulation during all tasks; that is, the strength of alpha-gamma comodulation was positively correlated with alpha power (Tort et al., 2008; Tort et al., 2013). These results are consistent with previous studies where stronger alpha-gamma comodulation occurred while alpha band activity increased (Osipova et al., 2008; Voytek et al., 2010; van Kerkoerle et al., 2014). Similar alpha-gamma comodulation was modulated by different reward conditions in the nucleus accumbens of humans as well (Cohen et al., 2009). Based on these previous studies and our results, we suggest that reduced dopamine release leads to a decrease in attention/motivation and manifested as an increased alpha power and increased alpha-gamma comodulation during nonrewarding trials in M1 cortex.

### Reward-Modulated Changes in Spike Field Coherence (SFC)

As shown in Figure 4, a larger subpopulation of reward-modulated M1 units had significantly higher phase synchronization as measured with SFC between alpha oscillations and neural spikes in nonrewarding trials than in rewarding trials for all tasks. This indicates that higher phase-synchronization is seen in the presence of stronger alpha oscillations. Our SFC results are consistent with a previous study (Haegens et al., 2011), which showed that an increase of alpha power was associated with an increase in alpha-band SFC in premotor, motor, and somatosensory regions during a discrimination task. This suggests that stronger alpha oscillatory activity in M1 may give rise to a suppression in neural spiking activity and modulates the inhibitory timing of neural spiking (Klimesch et al., 2007; Jensen and Mazaheri, 2010; Mazaheri and Jensen, 2010; Jensen et al., 2012; Klimesch, 2012). Thus, a possible explanation for higher phase synchronization in the alpha band between spikes and LFPs is that the lack of reward expectation leads to a decrease in dopamine associated motivation and/or attention during nonrewarding trials, which amplifies alpha-band LFP oscillations, and as a result alpha oscillations lead to the inhibition and timing of neural spiking. It should be kept in mind that these results were seen even when the NHPs had to perform manually or observe passively the reaching task or the force controlled grip force task, and thus they still needed to attend to the tasks to some extent, and thus some of these differences are likely due to decreased motivation.

### Reward-Modulated Changes in Neural Spike rate

As seen from the results in Figure 5, the neural firing rate of M1 units is modulated by cued reward expectation. These results include units that fire significantly lower or higher in the presence and expectation of reward. These observations are consistent with our previously reported findings (Marsh et al., 2015). It is interesting to see similar reward-related modulation in observational tasks and manual tasks, which indicate that these units might not code for movement alone but also for reward expectation (McNiel et al., 2016; Ramakrishnan et al., 2017; Tarigoppula et al., 2018). For instance, a large precue population (48.5%, 367 M1 units) in NHP A’s observational task is representative of the complete predictability of the reward schedule resulting from a sequenced presentation of rewarding and nonrewarding trials (Tarigoppula et al., 2018). A phenomenon which might partially explain similar population characteristics in observational tasks is the activity seen in mirror neurons, which are known to fire during the execution of a goal directed action as well as the observation of such an action (Tkach et al., 2008), and can even be modulated by subjective value (Caggiano et al., 2012). Therefore, this work, and previous work by our group and others, could be pointing to the fact that there are also mirror neurons modulated by value in M1 and S1 (Marsh et al., 2015; McNiel et al., 2016; Ramkumar et al., 2016; Ramakrishnan et al., 2017; An et al., 2018; Tarigoppula et al., 2018). More research is needed to further explain the detailed contributions mirror neurons have based on the results we have seen in observational tasks.

### Alpha Cycle Relation to Neural Spiking

In our results, we showed the possible existence of a relationship between the phase of alpha oscillations and neural firing rate (see Figure 6). We found that during nonrewarding trials for both manual and observational tasks in the contralateral M1 of NHPs A (for reaching tasks) and S (for grasping tasks) that the firing rate was high around the trough of the alpha cycle and low at the peak of the alpha cycle. These findings led to an initial conclusion that neural spiking was locked to the trough phase of the alpha band oscillations. The findings of the these results are consistent with a previous study (Haegens et al., 2011) where the firing rate was highest at the trough of the alpha cycle and lowest at the peak of its cycle. On the other hand, we saw slightly different results from our female NHPs in contralateral M1 units from NHPs P and AC8 (data not shown) as well as ipsilateral M1 units from NHP Z, where the firing rate was highest at the trough of the alpha cycle, but it was not lowest at the peak of the cycle. The reason for these results is not clear at the moment, but one possibility is that ipsilateral M1 units from NHP Z were less significantly modulated by a high expectation of reward than contralateral M1 units during arm reaching movement (Donchin et al., 1998; Cisek et al., 2003; Ganguly et al., 2009). Another interesting possibility is that this could be due to sex differences in dopamine receptor concentrations. Previous work has shown that female humans have much fewer dopamine receptors in the striatum as compared to males (Zaidi, 2010). These possibilities could have influenced the inhibition and timing of neural spiking with respect to the alpha cycle, although this is speculative.

The present findings seem to be consistent with the pulsed-inhibition theory (Klimesch et al., 2007; Jensen and Mazaheri, 2010; Mazaheri and Jensen, 2010; Haegens et al., 2011; Jensen et al., 2012; Klimesch, 2012) which is a mechanism through which alpha oscillations suppress neural spiking activity. All of our results are in line with the idea that alpha oscillations serve an inhibitory function and that stronger alpha power reduces task-relevant neural processing by decreasing firing rates (Haegens et al., 2011).

Future studies could be aimed at investigating inter-areal neural interactions (e.g., between M1, PMd, PMv, and S1) to investigate the bidirectional communication required for performing a variety of cognitive tasks (Pesaran et al., 2008; Tort et al., 2008; Terada et al., 2013; van Kerkoerle et al., 2014; Voloh et al., 2015; Arce-McShane et al., 2016). As mentioned above, the differences seen in rewarding versus nonrewarding trials through various measures have potential uses in improved learning for BMIs. Toward this goal, our lab recently showed that integrated features of PSD and SFC yielded near-perfect classification accuracy (An et al., 2018) between rewarding and nonrewarding trials during color-cued one-target center-out reaching tasks and thus can be used as a neural critic in autonomous BMI decoding (Bae et al., 2011; Sanchez et al., 2011; Tarigoppula et al., 2012; Marsh et al., 2015; An et al., 2018; Tarigoppula et al., 2018). Recently, we and others have shown that M1 directional tuning is also modulated by reward during manual (Ramakrishnan et al., 2017) and BMI control (Zhao et al., 2018), and that taking this into account could improve BMI control (Zhao et al., 2018). Further work is needed to incorporate our accurate classifier for an autonomously updating and accurate BMI system.

## Acknowledgments

This work was supported by NIH 1R01NS092894-01, NSF IIS-1527558, DARPA REPAIR Project N66001-10-C-2008, NYS Spinal Cord Injury Board Contracts C30600GG, C030838GG, and C32250GG.

